# Weak Relationship Between Divergence and Selection in a North American Toad Hybrid Zone

**DOI:** 10.1101/2024.08.09.606439

**Authors:** Kerry A. Cobb, Jamie R. Oaks

## Abstract

Hybrid zones present an excellent opportunity to study the evolution of reproductive isolation between divergent evolutionary lineages as generations of interbreeding and backcrossing produce many different recombinant genotypes. We can then observe patterns of genetic variation within a hybrid zone to understand how selection is acting on these recombinant genotypes. Using genome-wide sequence data, we characterized patterns of introgression within a putative hybrid zone between two species of North American toads (*A. americanus* and *A. terrestris*) to better understand reproductive isolation between these species. Both model based and non-parametric approaches to population structure inference showed that there is likely a substantial level of interbreeding and successful backcrossing at this hybrid zone with admixed individuals located quite far from the center of the hybrid zone. Bayesian genomic cline analysis revealed loci with extreme patterns of introgression relative to other loci which would be expected if these sites were linked to sites under negative selection in a hybrid genomic background. A site-based measure of genetic divergence was found to be weakly correlated with cline parameter estimates. We argue that this weak correlation is consistent with a history of secondary contact following a period of geographic isolation as many highly divergent sites do not have correspondingly high cline parameter estimates consistent with strong selection against introgression. Our findings substantiate previous claims of the existence of a hybrid zone between *A. americanus* and *A. terrestris* and highlight the potential for this hybrid zone to further our understanding of the evolution of reproductive incompatibly.

## 2 Introduction

The evolution of reproductive isolation between divergent lineages is a continuous process during which there may be ongoing gene flow or introgression via hybridization (Mallet, 2008; Wu, 2001). Introgression is possible because genetic barriers to introgression that accumulate within the genome are a property of genomic regions rather than a property of the entirety of the genome (Gompert, Parchman, et al., 2012; Wu, 2001). Natural hybridization between divergent lineages has become increasingly appreciated as a widespread phenomenon in recent years (Mallet, 2005; Moran et al., 2021). It is a phenomenon that can have important evolutionary consequences. Hybridization can be a source of adaptive variation (Hedrick, 2013). It can also introduce deleterious genetic load which persists long term within a population (Moran et al., 2021). Hybridization can potentially create conditions where selection favors the evolution of traits that enhance assortative mating and reduce the production of unfit hybrid offspring which drives further genetic divergence and reinforcement of reproductive barriers between lineages (Servedio & Noor, 2003). If hybrids do not suffer any negative fitness effects, hybridization could lead to the erosion of differences between divergent populations (Taylor et al., 2006), potentially resulting in populations that are genetically distinct from either parent species which can themselves eventually evolve reproductive isolation from the parent species (Moran et al., 2021).

Aside from having important evolutionary consequences which need to be understood, hybridization provides an excellent opportunity to investigate the processes that result in the evolution of reproductive incompatibility and divergence between evolutionary lineages. Hybrid zones are particularly suitable for this due to the production of a large number of recombinant genomes carrying many possible combinations of genomic elements from parent species resulting from generations of backcrossing (Rieseberg et al., 1999). Many generations of backcrossing and recombination make it possible to distinguish between the effects of closely linked genes (Rieseberg et al., 1999), and it is not feasible to achieve this experimentally in the vast majority of species (Rieseberg et al., 1999). Furthermore, the combination of genes produced are exposed to selection under natural conditions. This is important as the effect of hybrid incompatibilities can be dependent on environmental conditions and can only be fully understood in this context (Miller & Matute, 2016).

Despite being a fundamental evolutionary process, our understanding of evolution of reproductive incompatibility is far from complete (Butlin et al., 2011). Only a few loci, in a few species, have been pinpointed as the direct cause of reproductive incompatibility between species (Blackman, 2016; Nosil & Schluter, 2011). Consequently, our understanding of the processes that drive the evolution of loci resulting in reproductive incompatibility is limited (Butlin et al., 2011). Studies of introgression within hybrid zones have identified highly variable rates of introgression among loci (Barton & Hewitt, 1985; Gompert et al., 2017). This heterogeneity can arise via genetic drift occurring within hybrid zones, but will also be caused by differences among loci in the strength of selection against them in a hybrid genomic background (Barton & Hewitt, 1985; Gompert et al., 2017). It has also been observed that the levels of genetic divergence between species are highly variable across the genome (Nosil et al., 2009). Much of this heterogeneity is the result of divergent selection acting on each species independently (Nosil et al., 2009). Regions with particularly high levels of divergence between closely related species have been coined “genomic islands of divergence” (Wolf & Ellegren, 2017). It is assumed, particularly in the case of speciation with gene flow, that these genomic islands harbor genes that reduce the likelihood of successful interbreeding between species. When speciation occurs with gene flow, divergent selection can cause adaptive divergence in habitat use, phenology, or mating signals, and reduce the frequency or success of interspecific matings (Coyne & Orr, 2004). When species diverge in geographic isolation, divergent selection and reproductive isolation could be decoupled and reproductive isolation is not the result of direct selection against interspecific matings. Whether loci under divergent selection between two species also contribute to reproductive isolation has not been widely explored. A handful of studies have found evidence for a modest relationship between genetic divergence and selection against introgression (Gompert, Lucas, et al., 2012; Larson et al., 2013; Nikolakis et al., 2022; Parchman et al., 2013). How consistent and widespread this pattern is remains to be seen. At least one study published by Jahner et al. (2021) found no association.

In this study, we investigate hybridization between the American toad (*Anaxayrus americanus*) and Southern toad (*Anaxyrus terrestris*) at a suspected hybrid zone in the Southern United States to assess the extent of introgression between them and test for a relationship between introgression and genetic divergence. This suspected hybrid zone has not been investigated with genetic data previously, but it bears many hallmarks of a tension zone (Barton & Hewitt, 1985). Under the tension zone model of hybridization, species boundaries are maintained by a balance between dispersal and selection against individuals carrying incompatible hybrid genotypes (Barton & Hewitt, 1985). The ranges of *A. americanus* and *A. terrestris* abut with an abrupt transition and no apparent overlap along a long contact zone which ranges from Louisiana to Virginia. This contact zone closely corresponds with a prominent physiographic feature known as the “fall line” (Mount, 1975; Powell et al., 2016). The Fall line is the boundary between the Southern coastal plain to the South and the Appalachian Highlands to the North (Shankman & Hart, 2007). These regions differ in their underlying geology, topography, and elevation (Shankman & Hart, 2007). The distribution of *A. terrestris* is restricted to the coastal plain extending from the Mississippi River in the West to Virginia in the East (Fig. 1). The distribution of the American Toad encompasses nearly all of the Eastern North American with the exception of the Southern coastal plain (Fig. 1). Tension zones are expected to correspond with natural features that reduce dispersal or abundance and such a sudden transition is difficult to explain if not the result of the processes characteristic of tension zones (Barton, 1979). For there to be no mutually hospitable areas permitting some range overlap is implausible without there being an extreme level of competition or extreme degree of adaptation by each species to their respective environments. The two species only differ slightly in male advertisement call, morphological appearance, and the timing of their spawn (Cocroft & Ryan, 1995; Mount, 1975; Weatherby, 1982). Although there is some overlap in the spawning period and male Bufonidae are famously indiscriminate in their choice of mates (Ðorđević & Simović, 2014; Weatherby, 1982). They have also been shown to have a degree of reproductive compatibility through laboratory crossing experiments which produced viable *F*_2_ offspring (Blair, 1963). Analysis of morphological variation in central Alabama by Weatherby (1982) suggests there has been introgression between them.

**Figure 1.**
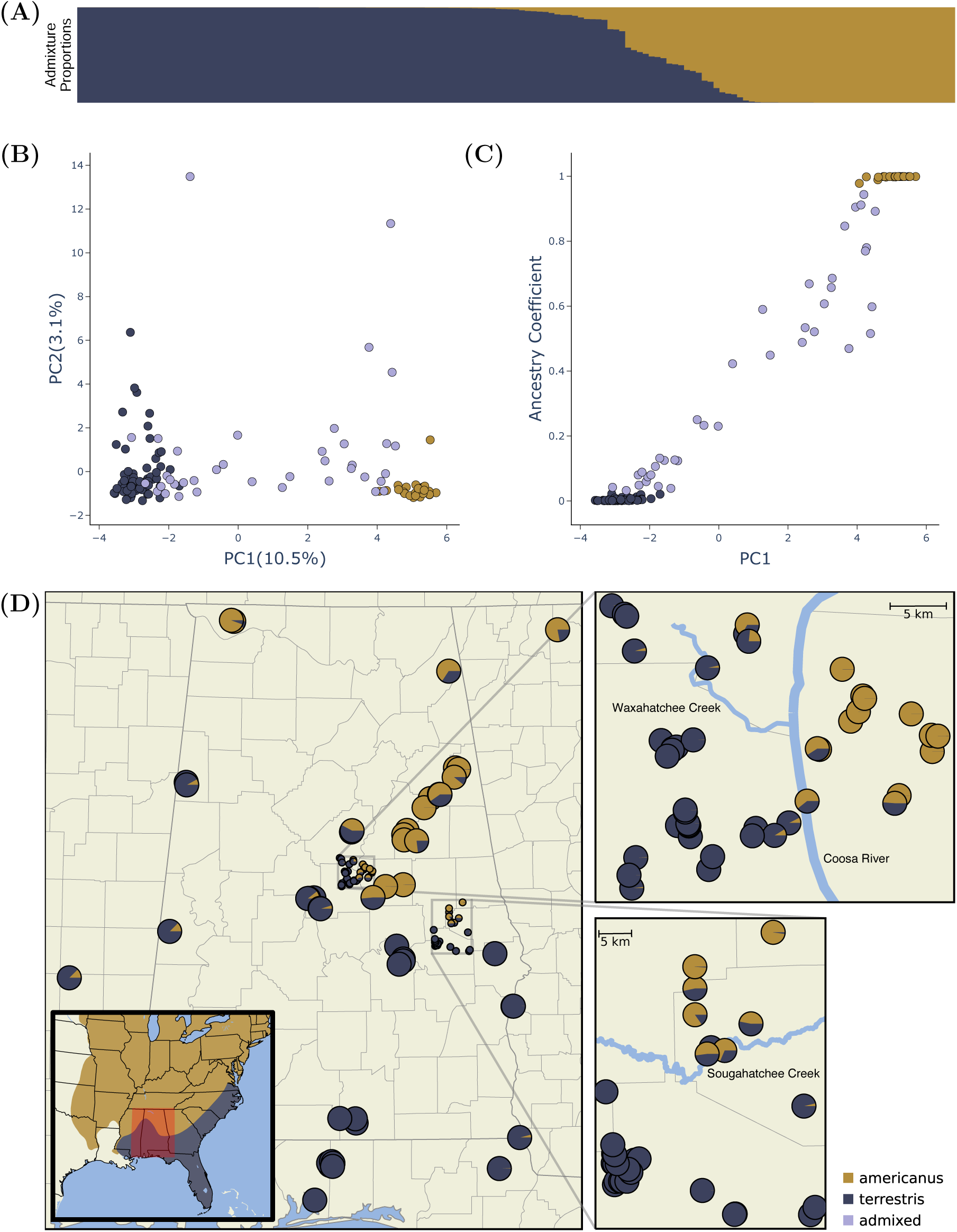
Genetic evidence of hybridization between *A. americanus* and *A. terrestris*. (A) Bar plot with the ancestry coefficients estimated with *STRUCTURE*. (B) Summary of poulation genetic structure based on the principal component axes one (PC1) and two (PC2). These axes explain 10.5% (PC1) and 3.1% (PC2) of the genetic variation among individuals. (C) Relationship between the first principal component axis and the admixture proportions estimated with *STRUCTURE*. (D) Sample map showing the sampling location and estimated ancestry coefficients of each sample. The inset map shows the approximate ranges of each species and the study area highlighted in red. Figure created using *POPHELPER* (Francis, 2017) and *Matplotlib* (Hunter, 2007)

The “true toads” in the family Bufonidae, to which *A. americanus* and *A. terrestris* belong, have been a prominent group of organisms in the literature on hybridization. W.F. Blair and colleagues performed a remarkable 1,934 separate experimental crosses to quantify the degree of reproductive incompatibility between species pairs within this family (Blair, 1972; Malone & Fontenot, 2008). These experiments demonstrated a high degree of compatibility between some closely related species pairs in which hybrids were capable of producing viable backcross or *F*_2_ hybrid offspring (Blair, 1963). Furthermore, numerous cases of natural hybridization among toad species have been reported with several apparent as well as indisputable hybrid zones (Colliard et al., 2010; Green, 1996; Van Riemsdijk et al., 2023; Weatherby, 1982). Despite the interest in and appreciation for hybridization in Bufonidae, only a small amount of work has been done to understand patterns of introgression within Bufonid hybrid zones. Just three hybrid zones have been investigated using genetic data. A clinal pattern of admixture at 26 allozyme loci has been shown within the *Anaxyrus americnaus* X *Anaxyrus hemiophrys* hybrid zone in Ontario, Canada(Green, 1983). Almost no admixture was detected at 7 microsatellite loci within the suspected *Bufo siculus* X *Bufo balearicus* hybrid zone in Sicily, Italy (Colliard et al., 2010). The most comprehensive study of introgression within a Bufonidae hybrid zone found significant levels of genome wide admixture, fitting a clinal pattern, at two separate transects at either end of the *Bufo bufo* x *Bufo spinosus* hybrid zone in Southern France (Van Riemsdijk et al., 2023).

The suspected *A. americanus*, *A. terrestris* hybrid zone has great potential to expand our understanding of the evolution of reproductive incompatibility. However, this will be dependent on the degree of ongoing introgression, if any, between these species. In this study, we use genome-wide sequence data to characterize patterns of introgression within the hybrid zone using model-based inference of admixture proportions, Bayesian genomic cline analysis, and estimates of parental population differentiation. With these approaches, we specifically address the following questions: 1) Is there evidence of ongoing hybridization and admixture between the two species, 2) Do any loci have outstanding patterns of introgression consistent with them being linked to reproductive incompatibility, and 3) Is there any relationship between patterns of introgression and levels of genetic differentiation between parental lineages?

## 3 Methods

### 3.1 Sampling and DNA Isolation

We collected genetic samples from *A. americanus* and *A. terrestris* by driving roads during rainy nights between 2017 and 2020 in a region of central Alabama where hybridization has previously been inferred from the presence of morphological intermediate individuals (Weatherby, 1982). For each individual collected, we euthanized it with immersion in buffered MS-222, preserved genetic samples of liver and/or toes in 100% ethanol, and preserved the rest of the specimen with 10% buffered formalin solution.

We euthanized individuals with immersion in buffered MS-222. We removed liver and/or toes and preserved them in 100% ethanol and fixed specimens with 10% Formalin solution. Genetic samples and formalin fixed specimens were deposited at the Auburn Museum of Natural History. Additional samples were also provided by museums (see Table S2).

We isolated DNA by lysing a small piece of liver or toe approximately the size of a grain of rice in a 300 *µ*L solution of 10mM Tris-HCL, 10mM EDTA, 1% SDS (w/v), and nuclease free water along with 6 mg Proteinase K and incubating for 4-16 hours at 55*^◦^*C. To purify the DNA and separate it from the lysis product, we mixed the lysis product with a 2X volume of SPRI bead solution containing 1 mM EDTA, 10 mM Tris-HCl, 1 M NaCl, 0.275% Tween-20 (v/v), 18% PEG 8000 (w/v), 2% Sera-Mag SpeedBeads (GE Healthcare PN 65152105050250) (v/v), and nuclease free water. We then incubated the samples at room temperature for 5 minutes, placed the beads on a magnetic rack, and discarded the supernatant once the beads had collected on the side of the tube. We then performed two ethanol washes by adding 1 mL of 70% ETOH to the beads while still placed in the magnet stand and allowing it to stand for 5 minutes before removing and discarding the ethanol. After removing all ethanol from the second wash, we removed the tube from the magnet stand and allowed the sample to dry for 1 minute before thoroughly mixing the beads with 100 *µ*L of TLE solution containing 10 mM Tris-HCL, 0.1 mM EDTA, and nuclease free water. After allowing the bead mixture to stand at room temperature for 5 minutes, we returned the beads to the magnet stand, collected the TLE solution, and discarded the beads. We quantified DNA in the TLE solution with a Qubit fluorometer (Life Technologies, USA) and diluted samples with additional TLE solution to bring the concentration to 20 ng/*µ*L.

### 3.2 RADseq Library Preparation

We prepared RADseq libraries using a the 2RAD approach developed by Bayona-Vásquez et al. (2019) with some minor modifications. On 96 well plates, we ligated 100 ng of sample DNA in 15 *µ*L of a solution with 1X CutSmart Buffer (New England Biolabs, USA; NEB), 10 units of XbaI, 10 units of EcoRI, 0.33 *µ*M XbaI compatible adapter, 0.33 *µ*M EcoRI compatible adapter, and nuclease free water with a 1 hour incubation at 37*^◦^*C. We then immediately added 5 *µ*L of a solution with 1X Ligase Buffer (NEB), 0.75 mM ATP (NEB), 100 units DNA Ligase (NEB), and nuclease free water and incubated at 22*^◦^*C for 20 min and 37*^◦^*C for 10 min for two cycles, followed by 80*^◦^*C for 20 min to stop enzyme activity. For each 96 well plate, we pooled 10 *µ*L of each sample and split this pool equally between two microcentrifuge tubes. we purified each pool of libraries with a 1X volume of SpeedBead solution followed by two ethanol washes as described in the previous section except that the DNA was resuspended in 25 *µ*L of TLE solution and combined the two pools of cleaned ligation product.

In order to be able to detect and remove PCR duplicates, we performed a single cycle of PCR with the iTru5-8N primer which adds a random 8 nucleotide barcode to each library construct. For each plate, we prepared four PCR reactions with a total volume of 50 *µ*L containing 1X Kapa Hifi Buffer (Kapa Biosystems, USA; Kapa), 0.3 *µ*M iTru5-8N Primer, 0.3 mM dNTP, 1 unit Kapa HiFi DNA Polymerase, 10 *µ*L of purified ligation product, and nuclease free water. We ran reactions through a single cycle of PCR on a thermocycler at 98*^◦^*C for 2 min, 60*^◦^*C for 30 s, and 72*^◦^*C for 5 min. We pooled all of the PCR products for a plate into a single tube and purified the libraries with a 2X volume of SpeedBead solution as described above and resuspended in 25 *µ*L TLE. We added the remaining adapter and index sequences which were unique to each plate with four PCR reactions with a total volume of 50 *µ*L containing 1X Kapa Hifi (Kapa), 0.3 *µ*M iTru7 Primer, 0.3 *µ*M P5 Primer, 0.3 mM dNTP, 1 unit of Kapa Hifi DNA Polymerase (Kapa), 10 *µ*L purified iTru5-8N PCR product, and nuclease free water. We ran reactions on a thermocycler with an initial denaturation at 98*^◦^*C for 2 min, followed by 6 cycles of 98*^◦^*C for 20 s, 60*^◦^*C for 15 s, 72*^◦^*C for 30 s and a final extension of 72*^◦^*C for 5 min. We pooled all of the PCR products for a plate into a single tube and purified the product with a 2X volume of SpeedBead solution as described above and resuspended in 45 *µ*L TLE.

We size selected the library DNA from each plate in the range of 450-650 base pairs using a BluePippin (Sage Science, USA) with a 1.5% dye free gel with internal R2 standards. To increase the final DNA concentrations, we prepared four PCR reactions for each plate with 1X Kapa Hifi (Kapa), 0.3 *µ*M P5 Primer, 0.3 *µ*M P7 Primer, 0.3 mM dNTP, 1 unit of Kapa HiFi DNA Polymerase (Kapa), 10 *µ*L size selected DNA, and nuclease free water and used the same thermocycling conditions as the previous (P5-iTru7) amplification. We pooled all of the PCR products for a plate into a single tube and purified the product with a 2X volume of SpeedBead solution as before and resuspended in 20 *µ*L TLE. We quantified the DNA concentration for each plate with a Qubit fluorometer (Life Technologies, USA) then pooled each plate in equimolar amounts relative to the number of samples on the plate and diluted the pooled DNA to 5 nM with TLE solution. The pooled libraries were pooled with other projects and sequenced on an Illumina HiSeqX by Novogene (China) to obtain paired-end, 150 base-pair sequences.

### 3.3 Data Processing

We demultiplexed the iTru7 indexes using the *process_radtags* command from *Stacks* v2.6.4 (Rochette et al., 2019) and allowed for two mismatches for rescuing reads. To remove PCR duplicates, we used the *clone_filter* command from *Stacks*. We demultiplexed inline sample barcodes, trimmed adapter sequence, and filtered reads with low quality scores as well as reads with any uncalled bases using the *process_radtags* command again and allowed for the rescue of restriction site sequence as well as barcodes with up to two mismatches. We built alignments from the processed reads using the *Stacks* pipeline. We allowed for 14 mismatches between alleles within, as well as between individuals (M and n parameters). This is equivalent to a sequence similarity threshold of 90% for the 140 bp length of reads post trimming. We also allowed for up to 7 gaps between alleles within and between individuals. We used the *populations* command from *Stacks* to filter loci missing in more than 5% of individuals, filter all sites with minor allele counts less than 3, filter any individuals with more than 90% missing loci, and to randomly sample a single SNP from each locus.

### 3.4 Genetic Clustering & Ancestry Proportions

To cluster individuals and characterize patterns of genetic differentiation and admixture between clusters, I used the Bayesian inference program *STRUCTURE* v2.3.4 (Pritchard et al., 2000) with *STRUCTURE* ’s admixture model which returns an estimate of ancestry proportions for each sample. To evaluate the assumption that samples are best modeled as inheriting their genetic variation from the two groups corresponding to the species identification made in the field, we ran *STRUCTURE* under four different models, each with a different number of assumed clusters of individuals (*K* parameter) ranging from 1 to 4. For each value of K, we ran 20 independent runs for 100,000 total steps with the first 50,000 as burnin. We used the R package *POPHELPER* v2.3.1 (Francis, 2017) to combine iterations for each value of *K* and to select the model producing the largest Δ*K* which is the the model that has the greatest increase in likelihood score from the model with one fewer populations as described by (Evanno et al., 2005). We also examined genetic clustering and evidence of admixture using a non-parametric approach with a principal component analysis (PCA) implemented in the R package *adegenet* v2.1.10 (Jombart, 2008). We visualized the relationship between the first principal component axis and the model based estimates of admixture proportion for each individual to check for agreement between the parametric *STRUCTURE* analysis and the non-parametric PCA analysis.

### 3.5 Genomic Cline Analysis

To investigate patterns of introgression across the hybrid zone we used the Bayesian genomic cline inference tool *BGC* v1.03 (Gompert & Buerkle, 2012) to infer parameters under a genomic cline model. A genomic cline model has two key parameters, denoted *α* and *β*, which describe introgression at each locus based on the estimated ancestry proportion or hybrid index of individuals. The *α* parameter is referred to as the cline center which is the increase (positive value) or decrease (negative value) in the probability of ancestry from one species for a given locus. The *β* parameter affects the cline rate which is the rate of transition from a low probability to a high probability of ancestry for one species at a given locus. We classified a sample as being admixed for the *BGC* analysis if it had an inferred admixture proportion of <95% for one species under the model with a *K* of two in the *STRUCTURE* analysis. We used *VCFtools* v0.1.17 to remove all non-biallelic sites from the the VCF file produced by the *populations* command in *Stacks*. We converted the VCF formatted data into the *BGC* format using *bgc_utils* v0.1.0, a *Python* package that we developed for this project (github.com/kerrycobb/bgc_utils). We ran *BGC* with 5 independent chains, each for 1,000,000 steps and sampled every 1000. We visualized MCMC output to confirm patterns consistent with the chains converging on a shared stationary distribution, discarded burnin samples, combined the independent chains, and identified outlier loci with *bgc_utils*.

The primary goal of *BGC* analysis is to identify loci which have exceptional patterns of introgression. These loci, or loci in close linkage to them, are expected to be enriched for genetic regions affected by selection due to reproductive incompatibility between the two species. We identified loci with exceptional patterns of introgression using two approaches described by Gompert and Buerkle (2011). (1) If locus-specific introgression differed from the genome-wide average, which we will refer to as “excess ancestry” following Gompert and Buerkle (2011). More specifically, we classified a locus as having excess ancestry if the 90% highest posterior density interval (HPDI) for the alpha or beta parameter did not cover zero. (2) If locus-specific introgression is statistically unlikely relative to the genome-wide distribution of locus-specific introgression, which we refer to as “outliers” following Gompert and Buerkle (2011). We classified a locus as an outlier if the median of the posterior sample for the *α* or *β* parameters for a locus was not contained in the interval from 0.05 to 0.95 of the cumulative probability density functions *Normal*(0*, τ_α_*) or *Normal*(0*, τ_β_*) respectively, where *τ_α_* and *τ_β_* are the median values from the posterior sample for the conditional random effect priors on *τ_α_* and *τ_β_*. These conditional priors describe the genome-wide variation of locus-specific *α* and *β*. We further classified outlier *α* parameter estimates for a locus based on whether the median of the posterior sample was positive or negative. Positive estimates of *α* mean there is a greater probability of *A. americanus* ancestry in individuals at the locus relative to their hybrid index, whereas negative estimates of *α* mean there is a greater probability of *A. terrestris* ancestry.

### 3.6 Genetic differentiation and Introgression

To test for a relationship between patterns of introgression and genetic divergence, we used *VCFtools* to calculate the Weir and Cockerham (1984) *F_ST_* between each species using only the samples inferred through the *STRUCTURE* analysis to have >95% ancestry for one species under the model with a *K* of two (Danecek et al., 2011). The Weir and Cockerham *F_ST_* is calculated per-site and we calculated the per-site *F_ST_* for the same sites as those used in the *BGC* analysis. To determine if patterns of introgression are correlated with population differentiation, we calculated Pearson’s correlation coefficients between *F_ST_* and the *α* and *β* parameters using *SciPy* with the default p-value calculation. To calculate the correlation coefficient, we used the absolute value of the median of the posterior sample for the *α* parameter and the median of the posterior sample for the *β* parameter. To further test for a relationship between population differentiation and *α*, we binned loci as positive alpha outliers, negative alpha outliers, and non-outliers. To test for differences in values of *F_ST_* among these three groups of loci, we used a Kruskal-Wallis test using *SciPy* v1.10.1 (Virtanen et al., 2020). To determine which groups differ significantly from each other, we used pairwise Mann-Whitney U tests with Bonferonni correction using *scikit-posthocs* (github.com/maximtrp/scikit-posthocs).

## 4 Results

### 4.1 Sampling and Data Processing

We prepared reduced-representation sequencing libraries from 173 samples collected for this study (Table S1) and 19 samples available from existing collections (Table S2)). The *Stacks* pipeline assembled reads into 432,336 loci with a mean length of 253.31 bp. Prior to filtering the mean coverage per sample was 32X. After filtering loci missing from greater than 5% of samples, filtering sites with minor allele counts less than 3, filtering individuals with greater than 90% missing loci, and randomly sampling a single SNP from each locus, 1194 sites remained and 43 samples were excluded from further analyses leaving a total of 149. For the included samples, 56 had been identified as most closely resembling *A. americanus* and 93 had been identified as most closely resembling *A. terrestris*.

### 4.2 Genetic Clustering & Ancestry Proportions

A visual inspection of the *STRUCTURE* results shows that each run with same value for *K* converged on very similar results (Fig. 1). The *STRUCTURE* model with the largest Δ*K* was the model with a *K* of two (Fig. 2). Furthermore, individuals are inferred as having ancestry derived largely from only two ancestral groups even for *K* values of three and four. For these values of *K*, only a small amount of ancestry is attributed to the third or fourth ancestral groups for any individual sample (Fig. 3). Using a 95% estimated ancestry proportion as a cutoff for considering individuals to have pure ancestry, 36 samples were classified as pure *A. americanus*, 75 as pure *A. terrestris*, and 38 as being admixed. The proportions of admixture among the samples shows a clear gradient between 0 and 1 which is consistent with many individuals being the product of advanced-generation hybrids beyond the *F*_1_ generation. The transition of admixture proportions from one species to the other with distance from the locations of pure individuals with proportions closest to 0.5 being found in the center of this transition (Fig. 1).

### 4.3 Patterns of Introgression

Visualization of the MCMC output with trace plots and histograms of each parameter indicated that each of the five chains run in *BGC* quickly converged on the same parameter space. We conservatively discarded the first 10% of samples as burnin. The median of the posterior sample for cline center parameter *α* ranged from -0.525-0.494 across loci. The cline shape parameter *β* was less variable and ranged from -0.158-0.220 across loci. We identified 16 loci with “excess ancestry” for the *α* parameter relative to the genome wide average; i.e., the 90% HDPI does not cover 0. Of these, the median of the posterior sample for 5 of these loci was negative and for 11 loci was positive. Negative values represent a greater probability of *A. americanus* ancestry at a locus relative to the individual’s hybrid index whereas positive values represent a greater probability of *A. terrestris* ancestry. We identified 116 loci as being “outliers” for the *α* parameter; i.e., they are statistically unlikely relative to the genome-wide distribution of locus-specific introgression. Of these, the median of the posterior sample for 24 of these loci was negative and for 92 loci it was positive (Fig. 2). All 16 of the loci identified as having “excess ancestry” for the *α* parameter relative to the genome-wide average were also identified as “outliers” relative to the genome-wide distribution of locus-specific introgression. We did not classify any loci as having “excess ancestry” or as being “outliers” with respect to the *β* parameter.

**Figure 2.**
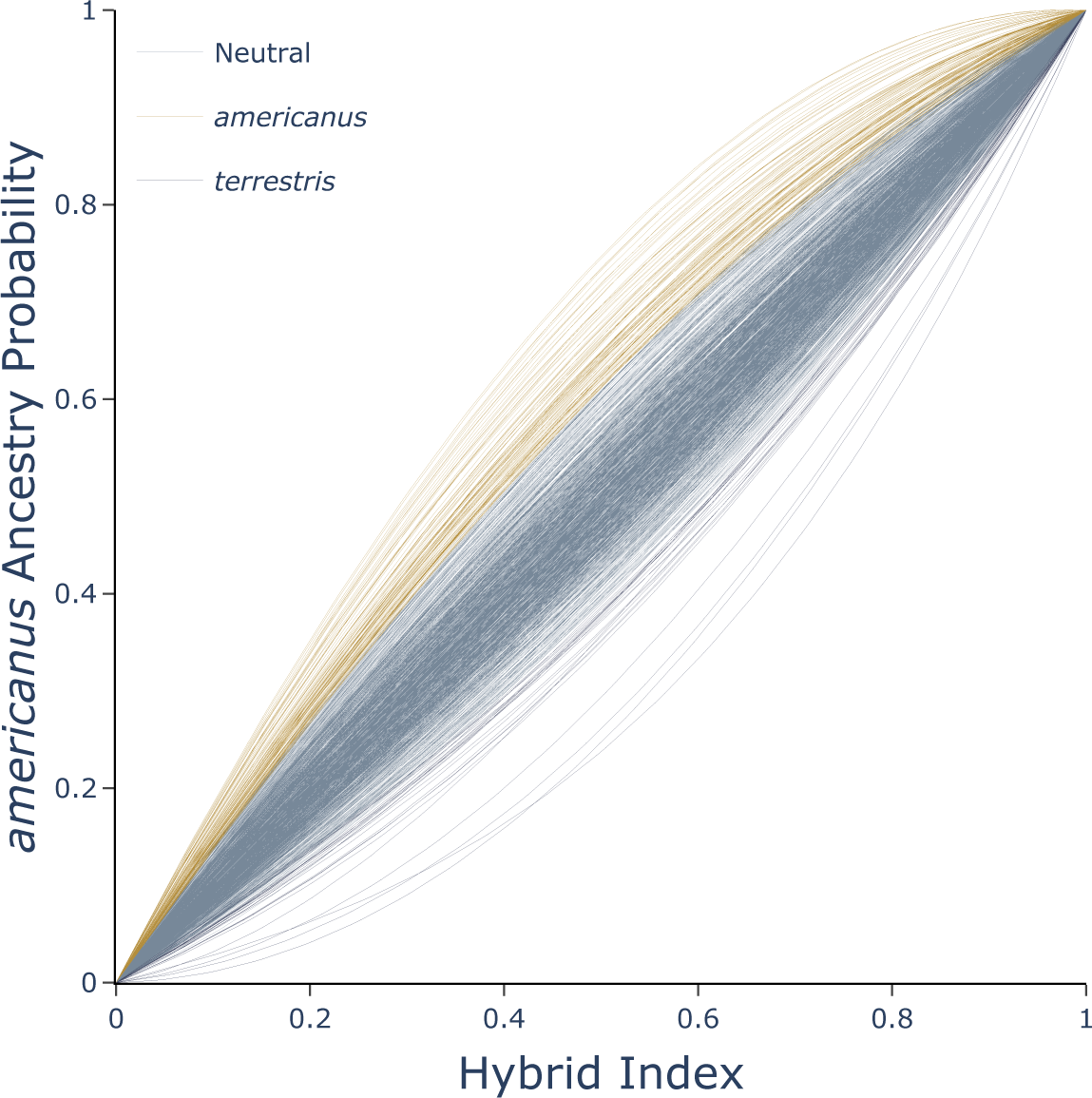
Shape of genomic clines estimated for each locus with *BGC*. Outliers are highlighted with yellow for loci that have greater *A. americanus* ancestry than expected and blue if loci have greater *A. terrestris* ancestry than expected.

### 4.4 Genomic Differentiation

Genetic differentiation between *A. americanus* and *A. terrestris* was highly variable among loci (Fig. 3). Locus-specific *F_ST_* between non-admixed *A. americanus* and *A. terrestris* had a mean of 0.07. *F_ST_*values for 249 loci were 0. Only a single locus had fixed differences between species with an *F_ST_* of 1.0. There is little apparent relationship between *α* or *β* and *F_ST_* except at the highest *α* and *β* estimates which have *F_ST_* estimates well above zero (Fig. 4). The Pearson correlation test estimates a weak correlation between *α* and *F_ST_* (r=0.29, p=1.62e-23) and between *β* and *F_ST_*(r=0.32, *p* = 8.28 *×* 10*^−^*^30^). The result of the Kruskal-Wallis test are consistent with there being significant differences between the *F_ST_* values of loci with outlier *α* estimates and non-outlier *α* estimates on average (*H* = 183.66, *p* = 1.32 *×* 10*^−^*^40^) (Fig. 3). The results of the post hoc pairwise Mann-Whitney tests are consistent with both categories of loci with outlier *α* estimates having greater *F_ST_* values on average than the non-outlier estimates of *α*. The difference between non-outlier loci and loci with greater probability of *A. americanus*ancestry was slightly higher (*U* = 88495, Bonferroni corrected *p* = 8.15 *×* 10*^−^*^38^) than the difference between non-outlier loci and loci with greater *A. terrestris* ancestry (*U* = 19519, Bonferroni corrected *p* = 2.45 *×* 10*^−^*^5^). Loci with greater probability of *A. americanus* ancestry had greater *F_ST_*on average than loci with greater probability of *A. terrestris* ancestry (*U* = 1741, Bonferroni corrected *P* = 4.32*e −* 05).

**Figure 3.**
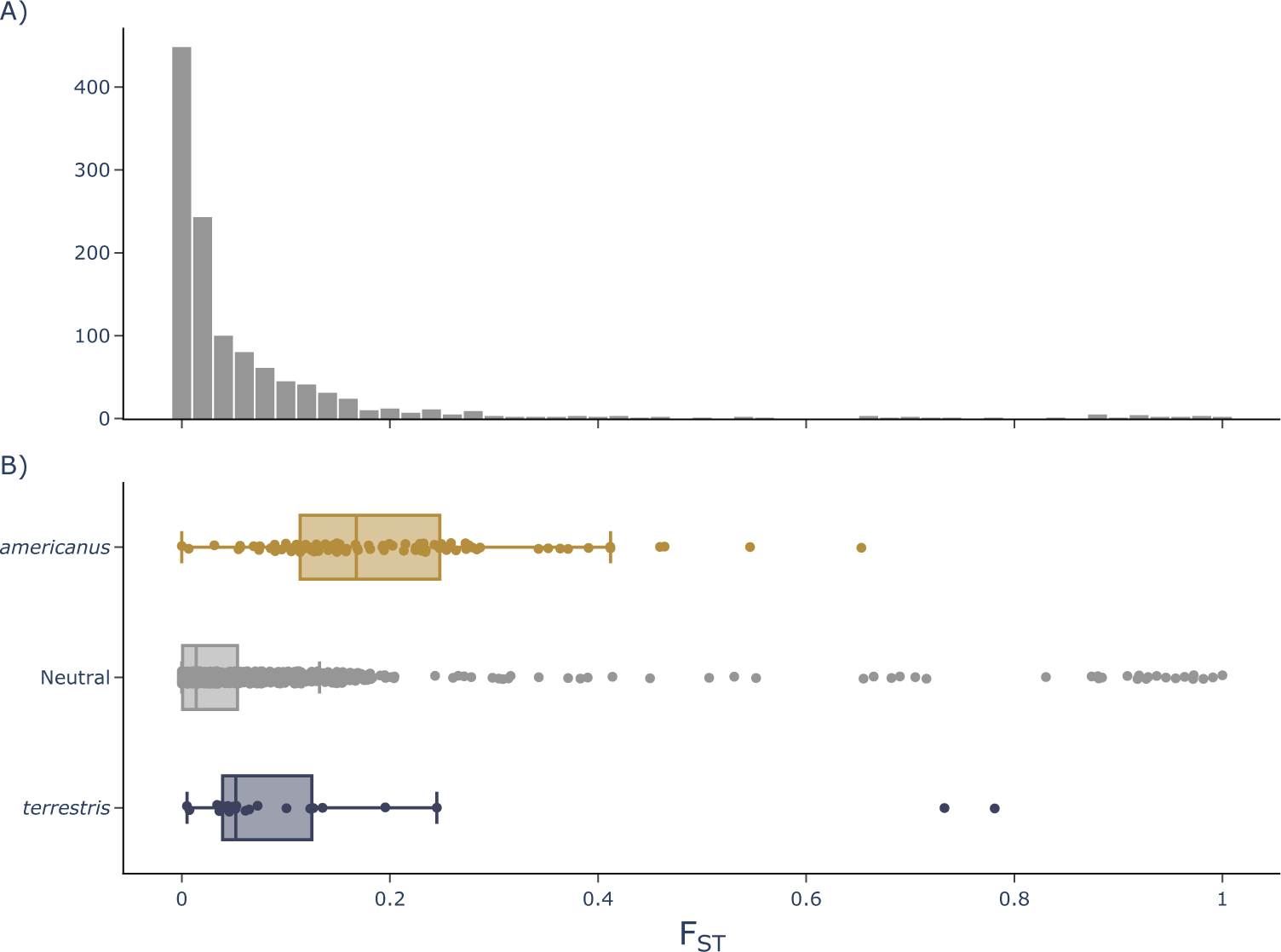
A) Distribution of per site *F_ST_* estimates. B) Box plots showing the distribution and mean of *F_ST_* for three categories of *α* estimates: outliers with greater than expected *A. americanus* ancestry (gold), outliers with greater than expected *A. terrestris* ancestry (violet), and non-outliers (gray).

**Figure 4.**
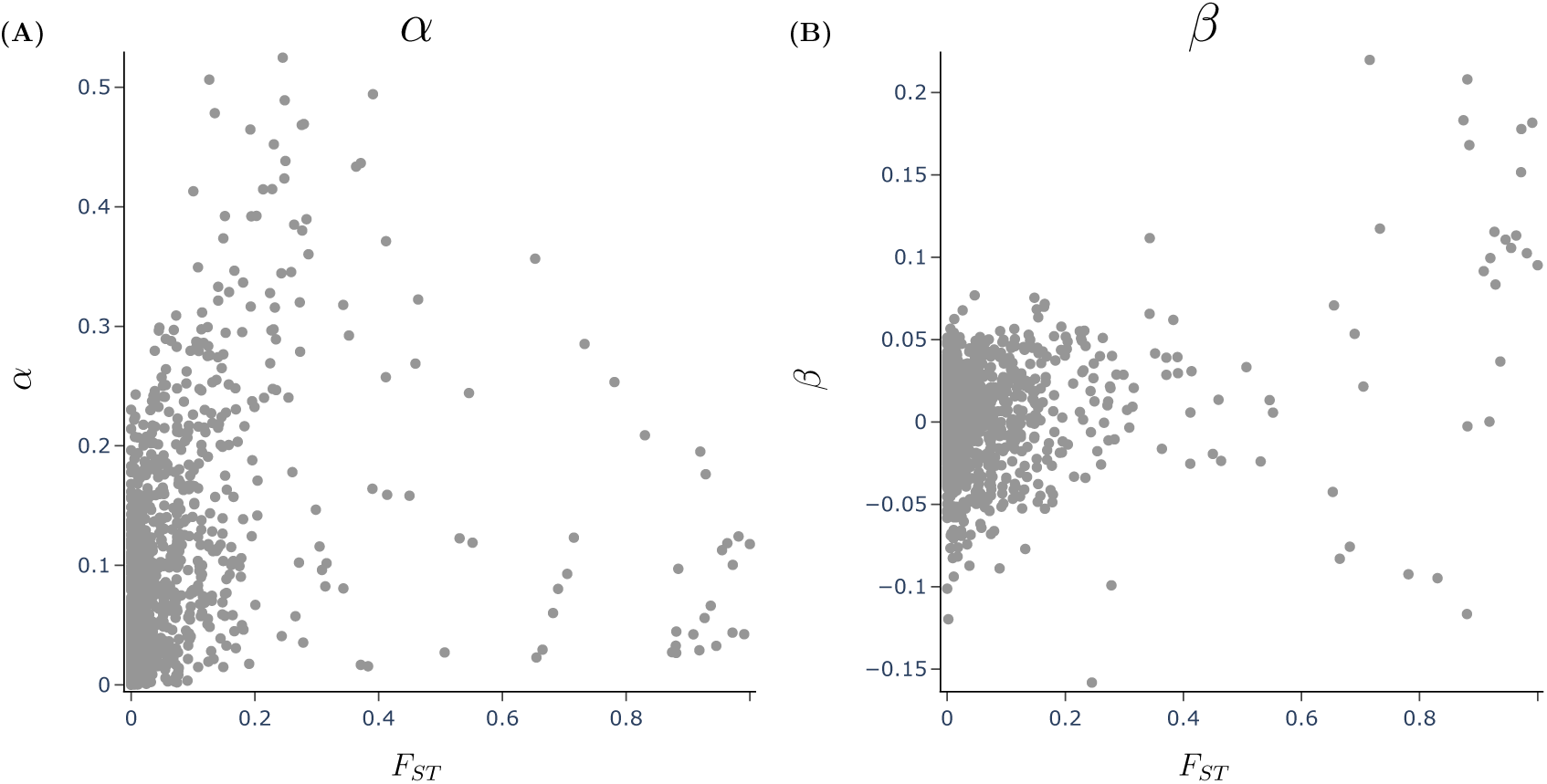
Relationship between genetic divergence measured with Weir and Cockerham, 1984 *F_ST_* and *BGC* cline parameters A) *α* and B) *β*.

## 5 Discussion

### 5.1 Evidence for ongoing hybridization

With the genome-wide sequence data obtained in this study, we find evidence of substantial gene flow across the hybrid zone of these two species. The *STRUCTURE* analysis inferred 38 out of 149 samples as having a proportion of ancestry of at least 5% of sites attributable to admixture (Fig. 1). The admixture proportions inferred in the *STRUCTURE* analysis range from 0.05-0.5 which is consistent with hybrids being viable, fertile, and capable of backcrossing over multiple generations (Fig. 1) (Slager et al., 2020). When backcrossing occurs over multiple generations in combination with migration of hybrid progeny and selection against introgressing alleles, a cline will form across the hybrid zone with introgressing alleles becoming more uncommon with distance from the cline edge (Barton & Hewitt, 1985). The results of the *STRUCTURE* analysis are largely consistent with this. Inferred admixture coefficients are highest at the center of the hybrid zone and decrease and approach zero with distance from the center (Fig. 1). Admixed samples were located quite far from the center of the hybrid zone. In fact samples with greater than 5% admixture proportions are located all the way at the Northeastern and Southwestern edges of the sampling area. The width of a hybrid zone is a product of the strength of selection for or against introgression and the average dispersal distance of individuals within their reproductive lifespan (Barton & Hewitt, 1985). Breden (1987) estimated that 27% of individual *A. fowleri* breed at non-natal breeding ponds with some dispersing more than 2 km. Female *A. americanus* have been shown to migrate more than 1 km between breeding sites and post-breeding locations (Forester et al., 2006). Invasive cane toads (*Rhinella marina*) in Australia are estimated to have expanded their range at a rate of 10-15 km per year shortly after their introduction although this rate slowed with time (Urban et al., 2008). The presence of samples with little to no admixture in close proximity to toads with high proportions of admixture shows that dispersal patterns may have an important roll in shaping the patterns of introgression in this hybrid zone. Individuals would be expected to appear more like their neighbors if dispersal rates and distances were very low and homogeneous. It is also likely that this hybrid zone may be more appropriately described as a mosaic hybrid zone rather than a more simple tension zone (Harrison, 1986). However, the sampling for this study is too sparse and irregular to definitively test this. Another possibility is that some of this inferred admixture is the result of a statistical artifact or due to error. *STRUCTURE* can only model admixture and not ancestral polymorphism which would be classified by the program as admixture (Pritchard et al., 2000). Some reassurance is provided by the result of the PCA which is largely consistent with the *STRUCTURE* results although it is possible that they could be affected by the same bias or error introduced in data collection and processing (Fig. 1).

The tension zone model of hybrid zones predicts that location of hybrid zones centers will be dependent on the effects of selection along with population density and natural dispersal barriers (Barton, 1979). The *STRUCTURE* results show that in two areas, there is a clear transition from samples with primarily *A. americanus*ancestry to samples with primarily *A. terrestris* ancestry corresponding with the locations of streams and rivers. In the Northern part of the sampling area, transitions occur at the Coosa River and at Waxahatchee Creek (Fig. 1). In the Southern part, they occur at Sougahatchee Creek (Fig. 1). Clearly these are not impassable boundaries as there has been introgression beyond them. However, they likely reduce dispersal and as a result the center of the hybrid zone is caught in this location as described by Barton (1979).

### 5.2 Variability of introgression

There are two primary parameters of interest in a genomic cline model that can be interpreted in the evolutionary context of hybrid zones. The *α* parameter specifies the center of the cline and is dependent on the increase or decrease in the probability of locus-specific ancestry from one of the parental populations. The *β* parameter specifies the rate of change in probability of ancestry along the genome-wide admixture gradient. Extreme estimates of these parameters may be associated with loci that cause reproductive incompatibility between hybridizing species. The Bayesian genomic cline analysis of the genome-wide data in this study yielded extreme estimates for *α* at some sites. Sites were classified as having extreme values in two ways. First, sites could be classified as having excess ancestry if the HDPI of *α* or *β* does not cover zero and is therefore extreme relative to the genome-wide average of cline parameter estimates. Second, sites could be classified as being outliers if they are extreme relative to the genome-wide distribution of locus-specific effects under the cline model. A greater number of sites qualified as outliers for estimates of *α* than qualified as having excess ancestry. There were 116 loci classified as outliers which make up 9.7% of the total number of sites. Of those, 16 were also classified as having excess ancestry making up 1.3% of all sites. This difference is consistent with other studies using both simulated and empirical data which typically find more outlier loci than excess ancestry loci (Gompert & Buerkle, 2012). Both of these methods can produce false positives as these extreme values can be produced solely by genetic drift rather than by by selection (Gompert & Buerkle, 2012). So not all sites with extreme estimates will be associated with incompatibility loci. The false positive rate is exacerbated when there are many loci with small effects on compatibility. However, these sites should be enriched for loci associated with modest to strong reproductive incompatibility and thus provide an upper estimate of the number of sites that are associated with these modest to strong barriers to gene flow (Gompert & Buerkle, 2012).

None of the estimates for *β* were classified as either outliers or as having excess ancestry. Simulations have demonstrated that the *α* parameter is more impacted by selection against hybrid genotypes than the *β* parameter (Gompert, Lucas, et al., 2012). Other studies have also found no extreme estimates of *β* (Gompert, Lucas, et al., 2012; Nikolakis et al., 2022). One possible interpretation of the absence of extreme values of *β* is that selection is only strong enough to have a significant impact on *α* but it is not strong enough to have a large impact on *β*. Unlike for *α*, there is not a strong relationship between locally positive selection favoring introgressed genotypes and *β* (Gompert, Lucas, et al., 2012). Therefore, some of the extreme values for *α* could be due to adaptive introgression which does not have much impact on estimates of *β*. This is plausible given the large extent of introgression we observed. Some of which is potentially due to adaptive introgression. It has been shown that there is a negative relationship between *β* and dispersal rate (Gompert, Lucas, et al., 2012). It is therefore also plausible that high dispersal rates, rather than selection is the cause of lower *β* values that do not reach the threshold to qualify as extreme.

Of the 9.7% of sites that qualified as *α* outliers, a substantially larger proportion had positive values which represent greater *A. americanus* ancestry than expected at those sites in admixed individuals. Negative *α* estimates represent a greater probability of *A. terrestris* ancestry at a site within admixed individuals. Sites with positive outlier estimates for *α* made up 7.7% of all sites whereas those with negative outlier estimates made up just 2%. This asymmetry suggests that introgression flows more in the direction of *A. americanus* than it does in the direction of *A. terrestris*. This result is consistent with a pattern evident upon visual inspection of the mapped *STRUCTURE* results. Samples collected from sites adjacent to sites with admixed samples appear to have a greater proportion of *A. terrestris* ancestry than *A. americanus* ancestry (Fig. 1). Taken together, these observations suggest that introgression at this hybrid zone is asymmetric (Yang et al., 2020). Asymmetries in introgression can arise for multiple reasons. There could be differences in mate choice which make females of one species more selective than females of the other (Baldassarre et al., 2014). There can also be species differences in dispersal tendencies. Reciprocal-cross differences in reproductive isolation, termed Darwin’s Corollary, are very common (Turelli & Moyle, 2007). If one of the sexes is more prone to dispersal, introgression will flow more freely in one direction that it would in the other. It is possible that this observation is just an artifact of sampling. Particularly if this is a highly mosaic hybrid zone. However, many more samples with primarily *A. terrestris* ancestry were collected than samples with primarily *A. americanus* ancestry.

### 5.3 Relationship between introgression and differentiation

Patterns of genetic differentiation and genomic introgression between *A. americanus* and *A. terrestris* are consistent with the hypothesis that regions of the genome experiencing divergent selection also affect hybrid fitness. As predicted, there is a positive association between locus-specific estimates of *F_ST_* and both the absolute value of the *α* and the *β* parameter estimates. Although this correlation supports the hypothesis that introgression outliers are linked to loci under selection, the association is only a modest one. Despite this, it is notable all of the outlier *α* estimates as well as the highest *β* estimates have *F_ST_* estimates well above zero. Whereas sites with lower *α* and *β* estimates span the entire range from zero to one. This is consistent with expectations of secondary contact where not all loci that have undergone genomic divergence will necessarily result in reproductive isolation. A tighter coupling of divergence and resistance to gene flow would be expected under a scenario of divergence with gene flow.

### 5.4 Conclusion

In conclusion, the genome-wide sequence data analysis conducted in this study has provided compelling evidence of significant gene flow across the hybrid zone of *A. americanus* and *A. terrestris*. The *STRUCTURE* analysis reveals that a substantial number of samples have ancestry proportions attributable to hybridization. These findings suggest that hybrids are not only viable and fertile but also capable of backcrossing over multiple generations. Furthermore, the spatial distribution of admixture coefficients suggests the formation of a cline, with the highest levels of admixture at the hybrid zone’s center gradually diminishing with distance. Patterns in the distribution of admixture coefficients suggest a potentially important roll for rivers as a partial barrier to dispersal. We found a weak relationship between loci with limited introgression and the degree of genetic divergence, measured with *F_ST_*. This study demonstrates that introgression between *A. americanus* and *A. terrestris* is ongoing and can serve as a guide for future studies which could leverage quantitative measures of prezygotic isolation such as calls or spawning period, reference genomes, or crossing experiments paired with cutting edge genomic tools to shed light on the process of speciation.

### 5.5 Data Availability

All raw sequence reads have been deposited in the NCBI Short Read Archive under BioProject PRJNA1108277. Sample accessions are listed in Table S1 and Table S1. Scripts used for analysis are available at github.com/kerrycobb/toad-evolution.

## 6 Acknowledgments

We thank Holden Smith and Charlotte Benedict for assistance in the field and laboratory. We also thank the University of Texas at El Paso Biodiversity Collections for the gifting of one sample used in this study. This work was completed in part with resources provided by the Auburn University Easley Cluster. This article is contribution number 964 of the Auburn University Museum of Natural History.

## Supplementary Material

**Figure S1.**
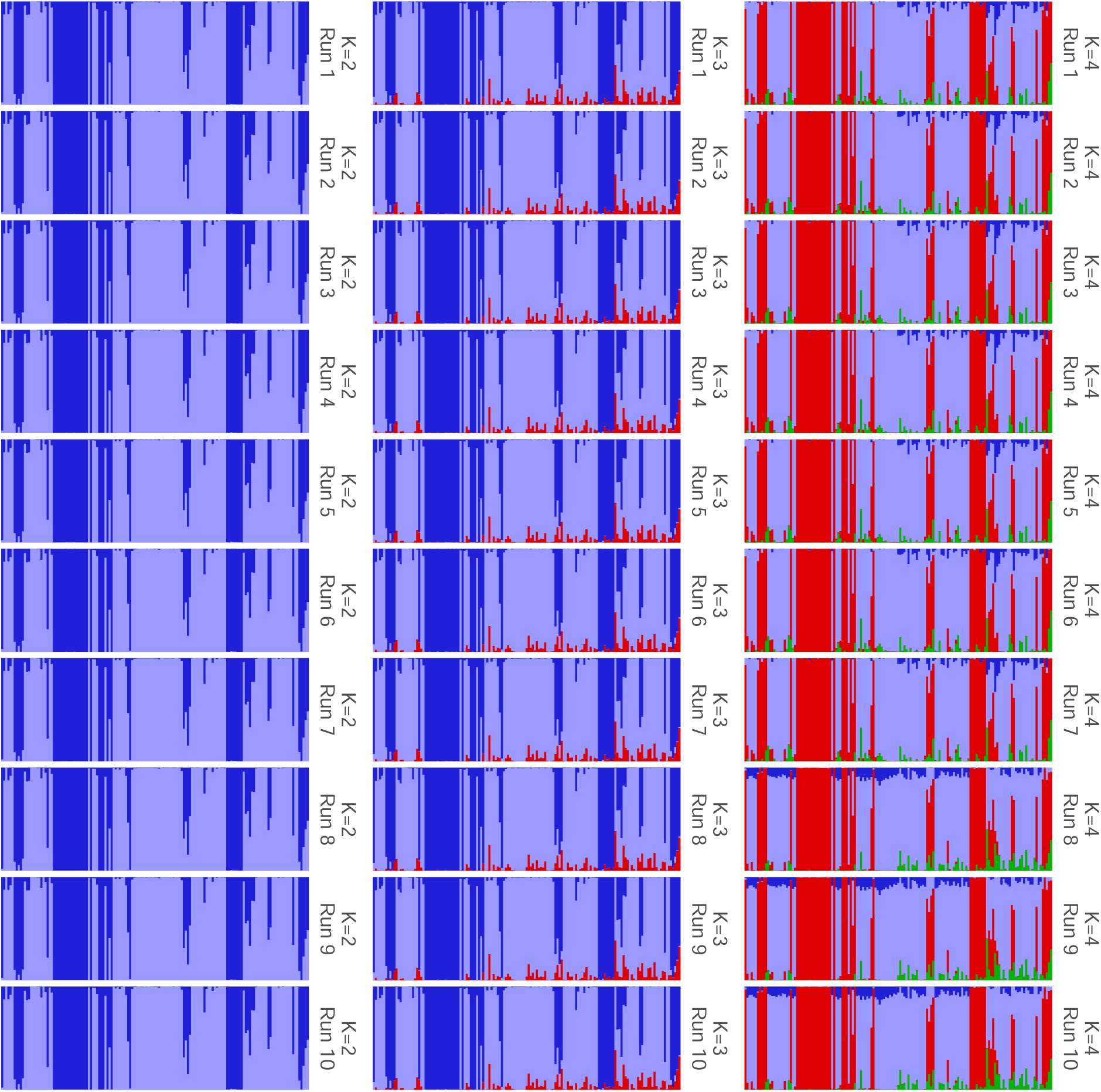
Results of each independent *STRUCTURE* run (rows) for each value of K (columns) showing convergence among runs with the same value for K. Plot was created with *POPHELPER* (Francis, 2017).

**Figure S2.**
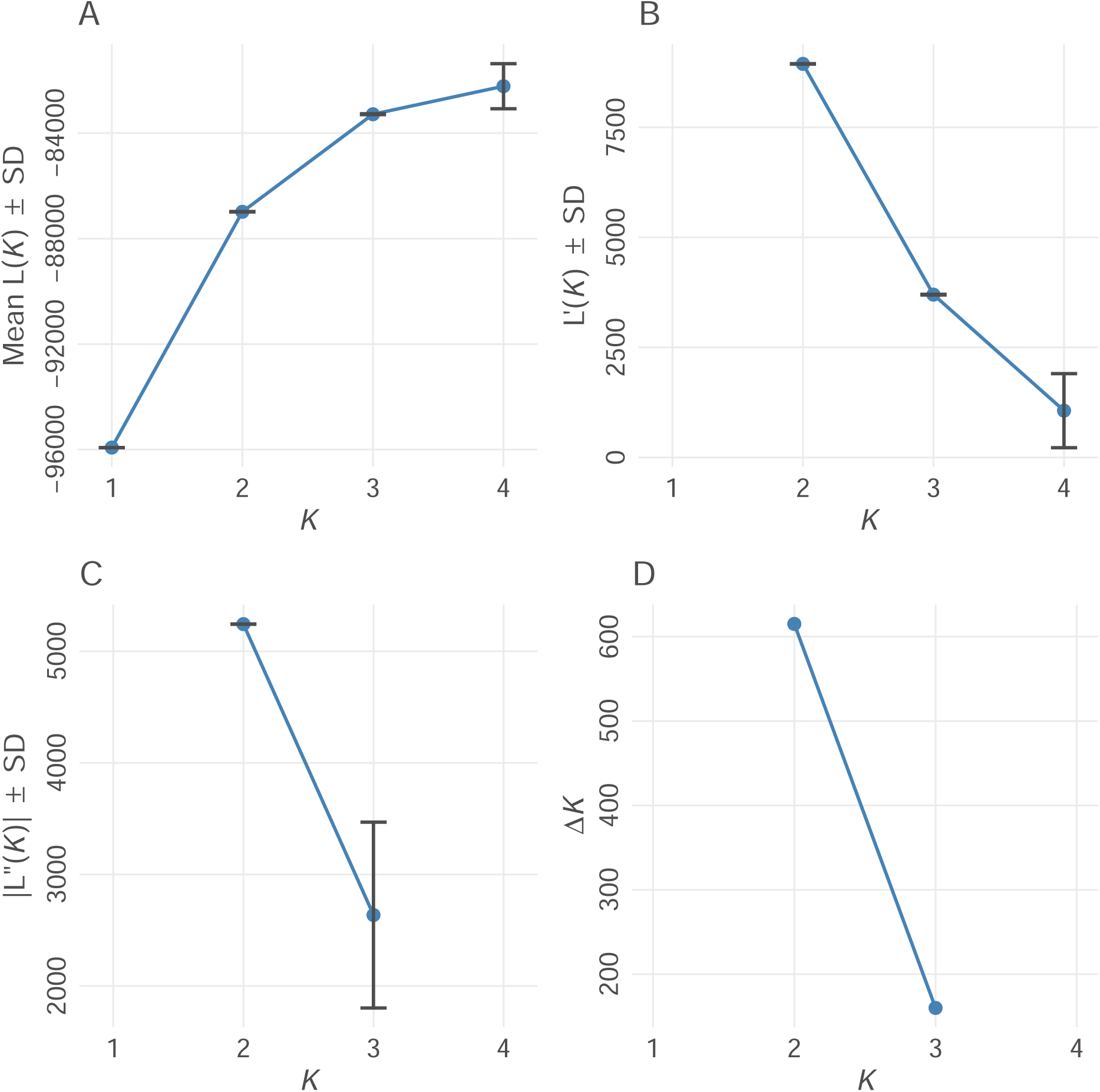
Evanno method for optimal value for K in *STRUCTURE* (Evanno et al., 2005). *K* refers to the number of populations for each of the different *STRUCTURE* models examined. (A) Mean estimated ln probability of data over 10 iterations for each value of *K ± SD*. (B) Rate of change of the likelihood distribution (mean *±SD*) (C) Absolute values of the second order rate of change of the likelihood distribution (mean *±SD*) (D) Δ*K*. The modal value of this distribution is considered the true value of *K* for the data. Plot created using *POPHELPER* (Francis, 2017).

**Figure S3.**
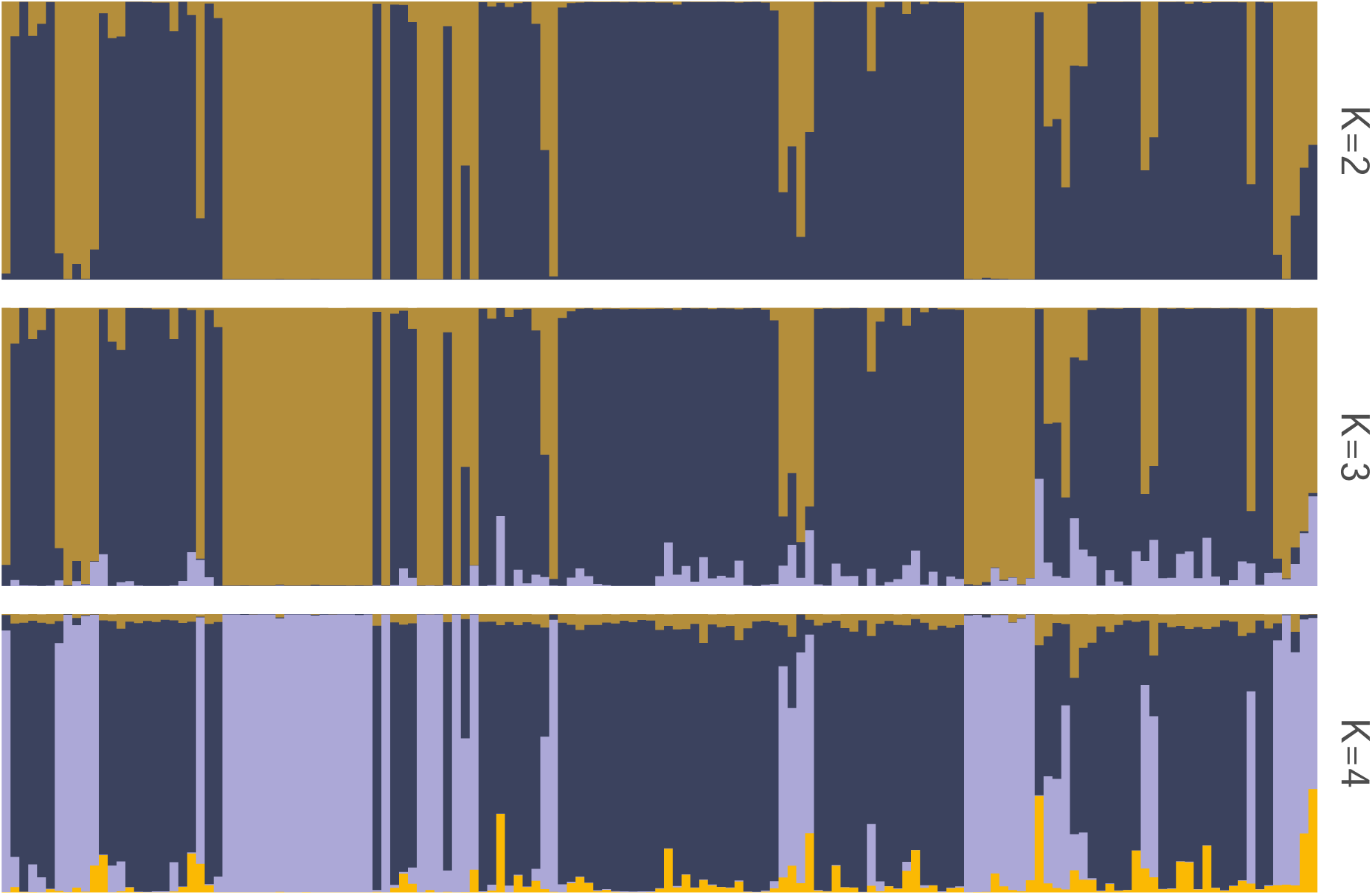
Summarized *STRUCTURE* results for each value of K. Ancestry proportions shown are the mean of ancestry proportions across all iterations. Sumarization and plotting done using *POPHELPER* (Francis, 2017).

**Table S1.** Samples collected for this study.

| ID | Voucher | SRA | Species | Passed Filter | Latitude | Longitude |
| --- | --- | --- | --- | --- | --- | --- |
| 046 | AUHT 7962 | SRR28909866 | *A. terrestris* | X | 30.54819 | -86.93067 |
| 047 | AUHT 7974 | SRR28909865 | *A. terrestris* | X | 32.81470 | -86.93968 |
| 048 | AUHT 7975 | SRR28909864 | *A. terrestris* | X | 32.81094 | -86.98967 |
| 049 | AUHT 7976 | SRR28909863 | *A. terrestris* | X | 32.80985 | -86.99795 |
| 050 | AUHT 7978 | SRR28909862 | *A. terrestris* |  | 32.82406 | -86.99314 |
| 051 | AUHT 7979 | SRR28909860 | *A. terrestris* |  | 32.82406 | -86.99314 |
| 052 | AUHT 7980 | SRR28909859 | *A. terrestris* |  | 32.80450 | -87.03078 |
| 053 | AUHT 7981 | SRR28909858 | *A. terrestris* |  | 32.76703 | -87.07073 |
| 054 | AUHT 7982 | SRR28909857 | *A. terrestris* |  | 32.76592 | -87.07184 |
| 055 | AUHT 7983 | SRR28909856 | *A. terrestris* |  | 32.78932 | -86.90850 |
| 056 | AUHT 7984 | SRR28909855 | *A. terrestris* | X | 32.73575 | -86.88149 |
| 057 | AUHT 7985 | SRR28909854 | *A. terrestris* | X | 32.73291 | -86.87707 |
| 058 | AUHT 7986 | SRR28909853 | *A. terrestris* |  | 32.74822 | -86.79806 |
| 059 | AUHT 7987 | SRR28909852 | *A. terrestris* |  | 32.78742 | -86.75847 |
| 060 | AUHT 7988 | SRR28909851 | *A. terrestris* |  | 32.78044 | -86.73877 |
| 069 | AUHT 8006 | SRR28909793 | *A. americanus* | X | 34.79963 | -84.57678 |
| 070 | AUHT 8007 | SRR28909792 | *A. terrestris* |  | 32.43478 | -85.64630 |
| 071 | AUHT 8010 | SRR28909790 | *A. terrestris* | X | 30.77430 | -85.22690 |
| 072 | AUHT 8011 | SRR28909789 | *A. terrestris* | X | 32.94778 | -86.63224 |
| 073 | AUHT 8012 | SRR28909788 | *A. terrestris* |  | 32.94970 | -86.52687 |
| 074 | AUHT 8013 | SRR28909787 | *A. terrestris* |  | 32.94970 | -86.52687 |
| 075 | AUHT 8014 | SRR28909786 | *A. americanus* | X | 33.00267 | -86.38960 |
| 076 | AUHT 8015 | SRR28909785 | *A. americanus* |  | 33.01205 | -86.47872 |
| 077 | AUHT 8016 | SRR28909784 | *A. americanus* |  | 33.04456 | -86.45547 |
| 078 | AUHT 8017 | SRR28909783 | *A. americanus* | X | 33.04456 | -86.45547 |
| 079 | AUHT 8018 | SRR28909782 | *A. americanus* | X | 33.04456 | -86.45547 |

Table S1 – continued from previous page
| ID | Voucher | SRA | Species | Passed Filter | Latitude | Longitude |
| --- | --- | --- | --- | --- | --- | --- |
| 080 | AUHT 8019 | SRR28909781 | *A. americanus* | X | 33.04456 | -86.45547 |
| 081 | AUHT 8020 | SRR28909779 | *A. americanus* | X | 33.04456 | -86.45547 |
| 082 | AUHT 8021 | SRR28909778 | *A. americanus* |  | 33.04456 | -86.45547 |
| 083 | AUHT 8022 | SRR28909964 | *A. americanus* | X | 33.04456 | -86.45547 |
| 084 | AUHT 8023 | SRR28909963 | *A. americanus* | X | 33.01484 | -86.39040 |
| 085 | AUHT 8025 | SRR28909962 | *A. americanus* | X | 33.01484 | -86.39040 |
| 086 | AUHT 8026 | SRR28909961 | *A. americanus* | X | 33.06472 | -86.47496 |
| 087 | AUHT 8027 | SRR28909960 | *A. americanus* | X | 33.06472 | -86.47496 |
| 088 | AUHT 8028 | SRR28909959 | *A. americanus* |  | 33.06472 | -86.47496 |
| 089 | AUHT 8029 | SRR28909958 | *A. americanus* | X | 33.06472 | -86.47496 |
| 090 | AUHT 8030 | SRR28909957 | *A. americanus* | X | 33.06472 | -86.47496 |
| 091 | AUHT 8031 | SRR28909955 | *A. americanus* | X | 33.06472 | -86.47496 |
| 092 | AUHT 8032 | SRR28909954 | *A. americanus* | X | 33.06472 | -86.47496 |
| 093 | AUHT 8033 | SRR28909953 | *A. americanus* | X | 33.06472 | -86.47496 |
| 094 | AUHT 8034 | SRR28909952 | *A. americanus* | X | 33.02572 | -86.46711 |
| 095 | AUHT 8035 | SRR28909951 | *A. americanus* | X | 33.02572 | -86.46711 |
| 096 | AUHT 8036 | SRR28909950 | *A. terrestris* | X | 32.92374 | -86.67199 |
| 097 | AUHT 8037 | SRR28909949 | *A. americanus* | X | 33.03283 | -86.45975 |
| 098 | AUHT 8038 | SRR28909948 | *A. terrestris* | X | 32.94544 | -86.55777 |
| 099 | AUHT 8039 | SRR28909947 | *A. terrestris* | X | 32.94947 | -86.52630 |
| 100 | AUHT 8040 | SRR28909946 | *A. terrestris* | X | 32.94947 | -86.52630 |
| 101 | AUHT 8041 | SRR28910057 | *A. americanus* | X | 33.04278 | -86.45377 |
| 102 | AUHT 8042 | SRR28910056 | *A. americanus* | X | 33.00464 | -86.49692 |
| 103 | AUHT 8043 | SRR28910055 | *A. americanus* | X | 33.01416 | -86.38417 |
| 104 | AUHT 8044 | SRR28910054 | *A. terrestris* | X | 32.94013 | -86.54004 |
| 105 | AUHT 8045 | SRR28910053 | *A. terrestris* |  | 32.94173 | -86.55787 |

Table S1 – continued from previous page
| ID | Voucher | SRA | Species | Passed Filter | Latitude | Longitude |
| --- | --- | --- | --- | --- | --- | --- |
| 106 | AUHT 8046 | SRR28910052 | *A. americanus* | X | 33.03099 | -86.40941 |
| 107 | AUHT 8047 | SRR28910051 | *A. americanus* | X | 33.00518 | -86.49895 |
| 108 | AUHT 8048 | SRR28910050 | *A. terrestris* |  | 32.95011 | -86.53723 |
| 109 | AUHT 8049 | SRR28910049 | *A. americanus* |  | 33.00528 | -86.38897 |
| 110 | AUHT 8050 | SRR28910048 | *A. americanus* |  | 33.01617 | -86.40318 |
| 111 | AUHT 8051 | SRR28910046 | *A. americanus* |  | 32.98218 | -86.40488 |
| 112 | AUHT 8052 | SRR28910045 | *A. americanus* | X | 32.96964 | -86.42137 |
| 113 | AUHT 8053 | SRR28910044 | *A. terrestris* |  | 32.97146 | -86.52901 |
| 114 | AUHT 8057 | SRR28910043 | *A. terrestris* | X | 32.44120 | -85.65386 |
| 115 | AUHT 8058 | SRR28910042 | *A. terrestris* |  | 32.85411 | -86.76619 |
| 116 | AUHT 8059 | SRR28910041 | *A. terrestris* | X | 32.90084 | -86.67587 |
| 117 | AUHT 8060 | SRR28910040 | *A. terrestris* | X | 32.91060 | -86.67850 |
| 118 | AUHT 8061 | SRR28910039 | *A. terrestris* |  | 32.91715 | -86.68208 |
| 119 | AUHT 8062 | SRR28910038 | *A. terrestris* |  | 32.92717 | -86.67407 |
| 120 | AUHT 8063 | SRR28910037 | *A. terrestris* |  | 32.97159 | -86.62516 |
| 121 | AUHT 8064 | SRR28910011 | *A. terrestris* |  | 33.00585 | -86.63703 |
| 122 | AUHT 8065 | SRR28910010 | *A. terrestris* |  | 33.00797 | -86.64210 |
| 123 | AUHT 8066 | SRR28910009 | *A. terrestris* |  | 33.00818 | -86.64333 |
| 124 | AUHT 8067 | SRR28910008 | *A. terrestris* |  | 33.01508 | -86.64937 |
| 125 | AUHT 8068 | SRR28910007 | *A. terrestris* |  | 33.02034 | -86.66651 |
| 126 | AUHT 8069 | SRR28910006 | *A. terrestris* | X | 33.01163 | -86.64759 |
| 127 | AUHT 8070 | SRR28910005 | *A. terrestris* | X | 33.00537 | -86.63652 |
| 128 | AUHT 8071 | SRR28910004 | *A. terrestris* | X | 33.00644 | -86.63368 |
| 129 | AUHT 8072 | SRR28910003 | *A. terrestris* | X | 33.00673 | -86.63316 |
| 131 | AUHT 8074 | SRR28910000 | *A. americanus* | X | 32.70224 | -85.66196 |
| 132 | AUHT 8075 | SRR28909999 | *A. americanus* | X | 32.73042 | -85.66173 |

Table S1 – continued from previous page
| ID | Voucher | SRA | Species | Passed Filter | Latitude | Longitude |
| --- | --- | --- | --- | --- | --- | --- |
| 133 | AUHT 8076 | SRR28909998 | *A. terrestris* | X | 32.62553 | -85.63684 |
| 134 | AUHT 8077 | SRR28909997 | *A. terrestris* | X | 32.41032 | -85.60107 |
| 135 | AUHT 8078 | SRR28909996 | *A. terrestris* | X | 32.57011 | -85.80888 |
| 136 | AUHT 8079 | SRR28909995 | *A. terrestris* | X | 32.47773 | -85.79824 |
| 137 | AUHT 8080 | SRR28909994 | *A. terrestris* | X | 32.47707 | -85.79577 |
| 138 | AUHT 8081 | SRR28909993 | *A. terrestris* | X | 32.48128 | -85.76354 |
| 139 | AUHT 8082 | SRR28909992 | *A. terrestris* | X | 32.48291 | -85.75622 |
| 140 | AUHT 8083 | SRR28909991 | *A. terrestris* | X | 32.45001 | -85.79652 |
| 141 | AUHT 8084 | SRR28909989 | *A. terrestris* | X | 32.45420 | -85.79408 |
| 142 | AUHT 8085 | SRR28909945 | *A. terrestris* | X | 32.45449 | -85.78664 |
| 143 | AUHT 8086 | SRR28909944 | *A. terrestris* | X | 32.45449 | -85.78664 |
| 144 | AUHT 8087 | SRR28909943 | *A. terrestris* | X | 32.45451 | -85.78416 |
| 145 | AUHT 8088 | SRR28909942 | *A. terrestris* | X | 32.45423 | -85.77634 |
| 146 | AUHT 8089 | SRR28909941 | *A. terrestris* | X | 32.45423 | -85.77634 |
| 147 | AUHT 8090 | SRR28909940 | *A. terrestris* | X | 32.46574 | -85.76977 |
| 148 | AUHT 8091 | SRR28909939 | *A. terrestris* | X | 32.46961 | -85.77369 |
| 149 | AUHT 8092 | SRR28909938 | *A. terrestris* | X | 32.47709 | -85.79175 |
| 151 | AUHT 8094 | SRR28909935 | *A. terrestris* | X | 32.47709 | -85.79175 |
| 152 | AUHT 8095 | SRR28909934 | *A. terrestris* | X | 32.49000 | -85.79741 |
| 153 | AUHT 8096 | SRR28909933 | *A. terrestris* | X | 32.40809 | -85.47857 |
| 154 | AUHT 8097 | SRR28909932 | *A. terrestris* | X | 32.41744 | -85.47117 |
| 155 | AUHT 8098 | SRR28909931 | *A. terrestris* | X | 32.35417 | -86.09838 |
| 156 | AUHT 8099 | SRR28909930 | *A. terrestris* | X | 32.33994 | -86.09946 |
| 157 | AUHT 8100 | SRR28909929 | *A. terrestris* | X | 32.31562 | -86.13789 |
| 160 | AUHT 8103 | SRR28909926 | *A. terrestris* | X | 33.06620 | -86.60328 |
| 161 | AUHT 8108 | SRR28909924 | *A. americanus* | X | 32.62171 | -85.61467 |

Table S1 – continued from previous page
| ID | Voucher | SRA | Species | Passed Filter | Latitude | Longitude |
| --- | --- | --- | --- | --- | --- | --- |
| 162 | AUHT 8109 | SRR28909923 | *A. americanus* | X | 32.61751 | -85.64335 |
| 165 | AUHT 8112 | SRR28909848 | *A. americanus* | X | 32.66836 | -85.66233 |
| 166 | AUHT 8113 | SRR28909847 | *A. americanus* | X | 32.65571 | -85.57134 |
| 170 | AUHT 8117 | SRR28909843 | *A. terrestris* | X | 32.38644 | -85.23561 |
| 171 | AUHT 8118 | SRR28909841 | *A. terrestris* | X | 32.38579 | -85.23565 |
| 172 | AUHT 8119 | SRR28909840 | *A. terrestris* | X | 32.38579 | -85.23565 |
| 173 | AUHT 8120 | SRR28909839 | *A. terrestris* | X | 32.38579 | -85.23565 |
| 174 | AUHT 8121 | SRR28909838 | *A. terrestris* | X | 32.38579 | -85.23565 |
| 176 | AUHT 8123 | SRR28909836 | *A. americanus* |  | 32.64548 | -85.55135 |
| 177 | AUHT 8124 | SRR28909835 | *A. terrestris* | X | 32.40976 | -85.60208 |
| 178 | AUHT 8125 | SRR28909834 | *A. terrestris* | X | 33.09152 | -86.56686 |
| 179 | AUHT 8126 | SRR28909833 | *A. terrestris* | X | 33.11298 | -86.69434 |
| 180 | AUHT 8127 | SRR28909832 | *A. terrestris* | X | 33.10659 | -86.68228 |
| 181 | AUHT 8128 | SRR28909830 | *A. terrestris* | X | 33.10509 | -86.68014 |
| 182 | AUHT 8129 | SRR28909829 | *A. terrestris* | X | 33.07896 | -86.67286 |
| 183 | AUHT 8130 | SRR28909828 | *A. terrestris* | X | 32.93933 | -86.62008 |
| 184 | AUHT 8131 | SRR28909827 | *A. terrestris* | X | 32.94745 | -86.62146 |
| 185 | AUHT 8132 | SRR28909826 | *A. terrestris* | X | 32.94829 | -86.62190 |
| 186 | AUHT 8133 | SRR28909988 | *A. terrestris* | X | 32.94929 | -86.62241 |
| 187 | AUHT 8134 | SRR28909987 | *A. terrestris* |  | 32.95077 | -86.62306 |
| 188 | AUHT 8135 | SRR28909986 | *A. terrestris* | X | 32.95794 | -86.62477 |
| 189 | AUHT 8136 | SRR28909985 | *A. terrestris* | X | 32.95940 | -86.62489 |
| 209 | AUHT 8141 | SRR28910034 | *A. terrestris* | X | 32.54852 | -85.48692 |
| 210 | AUHT 8142 | SRR28910033 | *A. americanus* | X | 33.30759 | -86.58201 |
| 211 | AUHT 8143 | SRR28910031 | *A. americanus* | X | 33.31685 | -86.57596 |
| 212 | AUHT 8144 | SRR28910030 | *A. americanus* | X | 33.09829 | -86.56529 |

Table S1 – continued from previous page
| ID | Voucher | SRA | Species | Passed Filter | Latitude | Longitude |
| --- | --- | --- | --- | --- | --- | --- |
| 213 | AUHT 8145 | SRR28910029 | *A. terrestris* | X | 33.08600 | -86.56394 |
| 214 | AUHT 8146 | SRR28910028 | *A. terrestris* | X | 33.08600 | -86.56394 |
| 215 | AUHT 8147 | SRR28910027 | *A. terrestris* |  | 33.01464 | -86.60995 |
| 216 | AUHT 8148 | SRR28910026 | *A. terrestris* | X | 33.01208 | -86.61707 |
| 217 | AUHT 8149 | SRR28910025 | *A. terrestris* | X | 33.00435 | -86.63710 |
| 218 | AUHT 8150 | SRR28910024 | *A. terrestris* | X | 32.99991 | -86.64181 |
| 219 | AUHT 8151 | SRR28910023 | *A. terrestris* |  | 32.99605 | -86.64526 |
| 220 | AUHT 8152 | SRR28910022 | *A. terrestris* |  | 33.01346 | -86.60960 |
| 221 | AUHT 8153 | SRR28910020 | *A. terrestris* | X | 32.91470 | -86.60270 |
| 222 | AUHT 8154 | SRR28910019 | *A. terrestris* | X | 32.92432 | -86.59895 |
| 223 | AUHT 8155 | SRR28910018 | *A. terrestris* | X | 32.93987 | -86.56113 |
| 224 | AUHT 8156 | SRR28910017 | *A. americanus* | X | 32.96579 | -86.50892 |
| 225 | AUHT 8157 | SRR28910016 | *A. americanus* | X | 32.96389 | -86.42549 |
| 226 | AUHT 8159 | SRR28910015 | *A. terrestris* |  | 32.53362 | -85.79839 |
| 227 | AUHT 8160 | SRR28910014 | *A. terrestris* | X | 32.48869 | -85.79555 |
| 228 | AUHT 8161 | SRR28910013 | *A. terrestris* | X | 32.50159 | -85.79860 |
| 230 | AUHT 8166 | SRR28909920 | *A. terrestris* | X | 30.80933 | -86.77686 |
| 231 | AUHT 8167 | SRR28909918 | *A. terrestris* | X | 30.80922 | -86.78994 |
| 232 | AUHT 8169 | SRR28909917 | *A. terrestris* | X | 30.80922 | -86.78994 |
| 233 | AUHT 8170 | SRR28909916 | *A. terrestris* | X | 30.80922 | -86.78994 |
| 234 | AUHT 8172 | SRR28909915 | *A. terrestris* | X | 30.82632 | -86.80258 |
| 235 | AUHT 8173 | SRR28909914 | *A. terrestris* | X | 30.83733 | -86.77630 |
| 236 | AUHT 8174 | SRR28909913 | *A. terrestris* | X | 30.82433 | -86.76284 |
| 237 | AUHT 8175 | SRR28909912 | *A. terrestris* | X | 30.80162 | -86.76659 |
| 240 | AUHT 8178 | SRR28909909 | *A. americanus* | X | 34.50446 | -85.63768 |
| 204 | AUHT 8181 | SRR28909967 | *A. americanus* |  | 34.21852 | -87.36662 |

Table S1 – continued from previous page
| ID | Voucher | SRA | Species | Passed Filter | Latitude | Longitude |
| --- | --- | --- | --- | --- | --- | --- |
| 190 | AUHT 8183 | SRR28909984 | *A. americanus* | X | 33.55274 | -85.82913 |
| 191 | AUHT 8186 | SRR28909982 | *A. americanus* | X | 33.48548 | -85.88857 |
| 192 | AUHT 8187 | SRR28909981 | *A. americanus* | X | 33.31649 | -86.05293 |
| 193 | AUHT 8188 | SRR28909980 | *A. americanus* | X | 33.28443 | -86.08443 |
| 194 | AUHT 8189 | SRR28909979 | *A. americanus* | X | 33.24576 | -86.08168 |
| 195 | AUHT 8190 | SRR28909978 | *A. americanus* |  | 32.91057 | -86.09272 |
| 197 | AUHT 8192 | SRR28909976 | *A. americanus* |  | 32.95104 | -86.14539 |
| 198 | AUHT 8193 | SRR28909975 | *A. americanus* | X | 32.89787 | -86.26061 |
| 199 | AUHT 8194 | SRR28909974 | *A. americanus* | X | 32.81642 | -86.38018 |
| 263 | AUHT 8198 | SRR28909811 | *A. terrestris* | X | 31.99016 | -85.07046 |
| 262 | AUHT 8200 | SRR28909812 | *A. terrestris* | X | 31.99042 | -85.07423 |
| 266 | AUHT 8201 | SRR28909808 | *A. americanus* | X | 33.23853 | -85.96270 |
| 265 | AUHT 8204 | SRR28909809 | *A. americanus* | X | 33.58435 | -85.74064 |
| 267 | AUHT 8205 | SRR28909807 | *A. americanus* | X | 33.58295 | -85.73539 |
| 264 | AUHT 8206 | SRR28909810 | *A. americanus* | X | 33.58295 | -85.73524 |
| 268 | AUHT 8207 | SRR28909806 | *A. americanus* | X | 32.81642 | -86.38018 |
| 261 | AUHT 8208 | SRR28909813 | *A. terrestris* |  | 31.10783 | -86.62247 |

**Table S2.** Samples loaned from museum collections.

| ID | Voucher | SRA | Species | Passed Filter | Latitude | Longitude |
| --- | --- | --- | --- | --- | --- | --- |
| 001 | AUHT 1975 | SRR28910060 | *A. americanus* | X | 32.77356 | -85.53325 |
| 002 | AUHT 2456 | SRR28910059 | *A. terrestris* | X | 32.19494 | -89.23629 |
| 005 | AUHT 2885 | SRR28909872 | *A. terrestris* | X | 32.45090 | -86.15934 |
| 007 | AUHT 3419 | SRR28909850 | *A. terrestris* | X | 33.67290 | -88.16068 |
| 008 | AUHT 3421 | SRR28909791 | *A. terrestris* | X | 33.65420 | -88.15580 |
| 009 | AUHT 3428 | SRR28909780 | *A. terrestris* | X | 31.12679 | -86.54755 |
| 010 | AUHT 3459 | SRR28909956 | *A. americanus* | X | 34.88028 | -87.71849 |
| 011 | AUHT 3460 | SRR28910058 | *A. americanus* | X | 33.78013 | -85.58421 |
| 012 | AUHT 3461 | SRR28910047 | *A. americanus* | X | 34.88779 | -87.74103 |
| 013 | AUHT 3462 | SRR28910012 | *A. americanus* | X | 33.77001 | -85.55434 |
| 014 | AUHT 3463 | SRR28910001 | *A. americanus* | X | 33.71125 | -85.59762 |
| 017 | AUHT 3813 | SRR28909925 | *A. terrestris* |  | 31.13854 | -86.53906 |
| 018 | AUHT 3833 | SRR28909842 | *A. terrestris* | X | 31.00422 | -85.03427 |
| 019 | AUHT 3997 | SRR28909831 | *A. terrestris* | X | 32.55607 | -88.29975 |
| 020 | AUHT 3998 | SRR28909983 | *A. terrestris* | X | 32.55607 | -88.29975 |
| 022 | AUHT 5276 | SRR28910032 | *A. terrestris* |  | 31.55613 | -86.82514 |
| 023 | AUHT 5277 | SRR28910021 | *A. terrestris* | X | 31.15830 | -86.55430 |
| 024 | AUHT 5278 | SRR28909919 | *A. terrestris* | X | 31.16105 | -86.69868 |
| 273 | UTEP 19947 | SRR28909896 | *A. terrestris* |  | 31.22432 | -88.77548 |

## Notes

### Competing Interest Statement

The authors have declared no competing interest.

### Summary of Updates

Changed the title and fixed some typos.

